# A common polymorphism in the druggable ion channel *PIEZO1* is associated with protection from severe malaria

**DOI:** 10.1101/691253

**Authors:** Christian N. Nguetse, Natasha Purington, Bikash Shakya, Emily R. Ebel, Peter G. Kremsner, Thirumalaisamy P. Velavan, Elizabeth S. Egan

**Affiliations:** Departments of Pediatrics and Microbiology & Immunology, Stanford University School of Medicine, Stanford, CA 94305; Quantitative Sciences Unit, Stanford University School of Medicine, Stanford, CA 94305; Department of Biology, Stanford University, Stanford, CA 94305; Institute of Tropical Medicine, University of Tübingen, Wilhelmstrasse 27, 72074 Tübingen, Germany; Centre de Recherches Médicales de Lambaréné, Albert Schweitzer Hospital, Lambaréné, Gabon; Vietnamese-German Center for Medical Research (VG-CARE), Hanoi, Vietnam; Duy Tan University, Da Nang, Vietnam

**Keywords:** malaria, *PIEZO1*, Africans, genetic association study

## Abstract

Malaria caused by the Apicomplexan parasite *Plasmodium falciparum* has served as a strong evolutionary force throughout human history, selecting for red blood cell polymorphisms that confer innate protection against severe disease. Recently, gain-of-function mutations in the mechanosensitive ion channel *PIEZO1* were shown to ameliorate *Plasmodium* parasite growth, blood-brain barrier dysfunction, and mortality in a mouse model of malaria. In humans, the gain-of-function allele *PIEZO1* E756del is highly prevalent and enriched in Africans, raising the possibility that it is under positive selection due to malaria. Here we used a case-control study design to test for an association between *PIEZO1* E756del and malaria severity among children in Gabon. We found that the E756del variant is strongly associated with protection against severe malaria in heterozygotes, independent of the protection conferred by the sickle cell trait (hemoglobin AS). *In vitro* experiments using donor red blood cells failed to find an effect of E756del on parasite growth, suggesting this variant confers a mild channel defect and/or that its protective effect may be mediated by other tissue types *in vivo*. Nonetheless, we show that Yoda1, a small molecule agonist of PIEZO1, has potent antimalarial activity in both E756del and wild-type red blood cells. Our findings demonstrate that *PIEZO1* is an important innate determinant of malaria susceptibility in humans and holds potential as druggable host target for malaria control.

## Introduction

Malaria infection due to *Plasmodium falciparum* is a major cause of childhood morbidity and mortality in endemic countries. The symptoms of the disease start when the parasite invades and replicates in red blood cells. Upon infection, there are three possible clinical outcomes that are influenced by the host, parasite, and environmental factors: uncomplicated malaria, severe malaria, or asymptomatic parasitemia. The development of asymptomatic parasitemia is largely influenced by adaptive immunity that results from repeated exposures, whereas it is estimated that ~25% of the clinical variation in malaria severity can be explained by innate genetic factors that act additively (MACKINNON *et al.* 2005). A variety of studies have shown that one of the strongest protective factors is heterozygosity for the hemoglobin S allele (HbAS), which leads to impaired parasite proliferation at low oxygen tension (PASVOL *et al.* 1978; JALLOW *et al.* 2009; TIMMANN *et al.* 2012; MALARIA GENOMIC EPIDEMIOLOGY *et al.* 2015; ARCHER *et al.* 2018). Many other candidate susceptibility loci reside in genes encoding membrane or structural proteins of the red blood cell (NGUETSE AND EGAN 2019). However, associations between established candidate loci and malaria severity explain only a fraction of the variance in clinical outcome, suggesting additional susceptibility factors have yet to be discovered (NDILA *et al.* 2018).

The mechanosensitive ion channel PIEZO1 was recently identified as a new candidate susceptibility factor for severe malaria (MA *et al.* 2018). PIEZO1 acts as a nonselective cation channel in a variety of tissues and has established roles in sensing blood flow through the vasculature, cell migration and differentiation, and red blood cell volume control (LI *et al.* 2014; PATHAK *et al.* 2014; RANADE *et al.* 2014; CAHALAN *et al.* 2015; WANG *et al.* 2016). Gain-of-function (GOF) mutations in *PIEZO1* underlie hereditary xerocytosis, a disorder characterized by red blood cell dehydration, reduced osmotic fragility and mild hemolytic anemia (ARCHER *et al.* 2014; GLOGOWSKA *et al.* 2017). Previous work has shown that *P. falciparum* parasites replicate poorly in severely dehydrated red blood cells, including those from patients with hereditary xerocytosis (TIFFERT *et al.* 2005). The demonstration that expression of a *PIEZO1*GOF allele in mice modeled the RBC abnormalities observed in hereditary xerocytosis, inhibited proliferation of *Plasmodium berghei* ANKA parasites, and protected the mice from cerebral malaria provided some *in vivo* evidence that GOF mutations in *PIEZO1* may influence susceptibility to severe malaria (MA *et al.* 2018).

Using a comparative genomics approach, a common *PIEZO1* polymorphism was identified in healthy individuals of African descent that may be under positive selection and act as a gain-of-function allele (MA *et al.* 2018). The mutation, E756del (rs572934641), is a 3 nucleotide deletion in a coding region of the gene that includes ~60 bp of short tandem repeats. PIEZO1 E756del was classified as a gain-of-function mutant based on studies in HEK cells, where it displayed a prolonged inactivation time constant after mechanical stimulation as compared to wild-type PIEZO1. Results from *in vitro* assays using the human malaria parasite *P. falciparum* suggested that parasite growth was impaired in RBCs from E756del heterozygotes as compared to wild-type RBCs (MA *et al.* 2018). Together, these findings raise the intriguing possibility that the *PIEZO1* E756del allele may be protective against severe malaria in humans.

Here, we screened a large case-control study group from Gabon for the *PIEZO1* E756del mutation and assessed its impact on the risk of severe malaria. We found that E756del was associated with protection from severe malaria in heterozygotes, suggesting that natural genetic variation in *PIEZO1* is an innate determinant of malaria susceptibility in humans. This association was independent of hemoglobin AS, which appears to have an epistatic interaction with *PIEZO1* E756del. While wild-type and E756del red blood cells supported the intracellular growth of *P. falciparum* equally well, we found that chemical upregulation of PIEZO1 inhibited *P. falciparum* replication. PIEZO1 is a druggable ion channel that may be under selective pressure to protect against severe malaria, suggesting that it holds potential as a target for a novel, host-directed therapy for this ancient disease.

## Methods

### Genetic Association study design and participants

Using a case-control approach, we assembled 542 samples from children aged 4-140 months that had been obtained in three previously published studies conducted in Lambaréné and Libreville, Gabon (KUN *et al.* 1998; KALMBACH *et al.* 2006; KREMSNER *et al.* 2009). After excluding samples for missing parameters, the analytic cohort consisted of 446 samples, of which 193 were controls with mild malaria and 253 were cases with severe malaria. All cases presented with microscopically confirmed *P. falciparum* parasitemia, signs and symptoms of severe malaria and no evidence of other severe diseases. Severe malaria was defined as severe anemia (hemoglobin <50 g/l) and/or hyperparasitemia (>250,000 parasites/μl, corresponding to >10% infected erythrocytes), a Blantyre coma score ≤2 and/or other facultative signs of severe malaria such as cerebral malaria, convulsions, hypoglycemia, and respiratory distress. The control group was those with mild malaria coming from the same geographical area as the cases. Mild malaria was defined as parasitemia 1000–50,000/μl on admission, no schizontemia, circulating leukocytes containing malarial pigment <50/μl, not homozygous for hemoglobin S, hemoglobin >80 g/l, platelets >50/nl, leukocytes <12/nl, lactate <3 mmol/l, and blood glucose >50 mg/dl. Written informed consent for each child was provided by the parents/guardians before enrollment.

### Mutation screening

*PIEZO1* exon 17 was amplified by PCR using the following primers: 5′-CAGGCAGGATGCAGTGAGTG-3′ (forward) and 5′-GGACATGGCACAGCAGACTG-3′ (reverse). Amplification reactions were performed in one batch in the same laboratory. The thermal conditions were an initial denaturation (95°C, 3 min) followed by 35 cycles of 95°C for 30 s, 65°C for 30 s and 72°C for 1 min. The PCR was completed with a final extension step of 72°C for 5 min. PCR products were visualized through electrophoresis on a 1·2% agarose gel stained with SYBR Safe DNA Gel Stain (Thermo Fisher Scientific, Waltham, MA, USA). Subsequently, the PCR products were purified with QIAquick PCR purification kit (Qiagen, Hilden, Germany) and directly used as templates for DNA sequencing (Quintara Biosciences, San Francisco, CA, USA). Mutations were identified by aligning the sequences with the *PIEZO1* reference sequence (NG_042229.1) using the Geneious 10.2.3 software (Auckland, New Zealand) and visually reconfirmed from their electropherograms. All samples were also genotyped for the HbS polymorphism (rs334) in the *HBB* gene (NG_059281.1). Briefly, we amplified exon 1 of the *HBB* gene using the primer pairs: 5′-AGTCAGGGCAGAGCCATCTA-3′ (forward) and 5′-GTCTCCACATGCCCAGTTTC-3′ (reverse). The PCR conditions were: initial denaturation (95°C, 3 min) followed by 35 cycles of 95°C for 30 s, 64°C for 30 s and 72°C for 1 min. The amplification was completed with a final extension step of 72°C for 5 min. The sequencing and variant detection were performed as described above.

### Statistical analysis

Participants with HbSS or HbSC hemoglobin genotypes were excluded from all analyses due to the severity of their underlying hematologic diseases. Additionally, all primary analyses were conducted only on participants without missing observations. Deviation from Hardy-Weinberg equilibrium was assessed using Chi-square analysis. Descriptive statistics were generated to evaluate the distributions of key characteristics by malaria severity status. The Wilcoxon rank sum test and Fisher’s exact test were used to determine if there were significant differences in malaria severity status in continuous and categorical characteristics, respectively.

The primary outcome was an indicator for whether a patient had mild or severe malaria. A main effects logistic regression model was fit to malaria severity status as a function of *PIEZO1* E756del genotype (normal [WT/WT=reference level], heterozygous [WT/DEL], and homozygous [DEL/DEL]), hemoglobin S genotype (normal HbAA, sickle cell trait HbAS), age in months, sex, and the study from which the participant information was extracted. Odds ratios (ORs) and 95% confidence intervals (CIs) were extracted from the logistic regression model. The estimated probability of severe malaria was extracted from the model for each participant and pairwise comparisons between the *PIEZO1* E756del variants and between hemoglobin S genotypes were presented. P-values for the pairwise differences in *PIEZO1* E756del were calculated from Tukey’s HSD (honestly significant difference) post hoc test. The Hosmer-Lemeshow goodness of fit was used to evaluate model fit using 10 bins.

In order to assess whether HbAS confounds the association between *PIEZO1* E756del and malaria severity status, an additional model was fit to malaria severity status as a function of *PIEZO1* E756del, HbAS, age, sex, study, and an interaction term between HbAS and *PIEZO1* E756del. Due to convergence issues based on small sample size, a Bayesian logistic regression model was used (GELMAN ANDREW AND YU-SUNG SU 2018). Similar methods to those used in the main effects model were implemented. Unadjusted logistic models were also fit to assess the association between seven additional *PIEZO1* polymorphisms near E756del and severe malaria.

Classification trees (CART) were implemented to determine whether a synergistic effect of any *PIEZO1* polymorphisms existed in order to predict malaria severity status. CART is a tree-based learning technique for classifying observations, which yields a decision rule by partitioning the data into subsets based on variables entered into the algorithm through the minimization of an error function. Two trees were grown to a maximum depth of 3 levels and minimal node size of 4 using the ctree function in the party package in R (HOTHORN *et al.* 2006), and included the following features, respectively: 1) all identified *PIEZO1* mutations, HbAS, age, and sex; 2) all identified *PIEZO1* mutations excluding *PIEZO1* E756del, HbAS, age, and sex. To ensure the validity of these models, we conducted a 5 times repeated 10-fold cross-validation (CV) for both CARTs. The CV area under the curve (AUC) and 95% CIs were reported for each tree (KOHAVI 1995). Spearman’s rank correlation test was also used in order to assess the genetic association (linkage) between *PIEZO1* E756del and the other *PIEZO1* mutations.

As a sensitivity analysis, missing values in age and sex were imputed by chained equations with predictive mean matching in order to assess the robustness of the two *PIEZO1* E756del logistic regression results (VAN BUUREN AND GROOTHUIS-OUDSHOORN 2011).

A p-value < 0·05 was considered statistically significant. All analyses were conducted using R software v3.5.2 (DEVELOPMENT CORE TEAM 2013).

### Blood sample collection and genotyping for in vitro parasite growth assays

Subjects who self-identified as African or African-American were recruited to donate blood at the Stanford Clinical Translational Research Unit. All participants and/or their parents gave informed consent according to a protocol approved by the Stanford University IRB (#40479). Whole blood samples were drawn into CPDA tubes and processed within 48 hours to remove serum and separate the buffy coat and red blood cells. Red blood cells were washed and stored in RPMI-1640 medium (Sigma) supplemented with 25 mM HEPES, 50 mg/L hypoxanthine, 2.42 mM sodium bicarbonate media at 4°C for up to 48 hours, and then cryopreserved in human AB+ serum and glycerol at −80°C. Genomic DNA was isolated from buffy coats using a DNeasy Blood and Tissue Kit (Qiagen). *PIEZO1* exon 17 was amplified by PCR of genomic DNA using the primers described above and Q5 polymerase in the following reaction conditions: 98°C for 30 seconds initial denaturation followed by 35 cycles of 98°C for 10 seconds, 70°C for 20 seconds, and 72°C for 20 seconds, followed by a 2 minute final extension at 72°C. PCR products were visualized on an agarose gel and sent for Sanger sequencing at Elim Bio (Hayward, CA). The *PIEZO1* E756 genotypes were manually determined from chromatograms examined using FinchTV software (Geospiza Inc). All genotypes were assessed by two independent researchers.

### *P. falciparum* culture and growth assays

*P. falciparum* strain 3D7 is a laboratory-adapted strain that was obtained from the Walter and Eliza Hall Institute (Melbourne, Australia) and was routinely cultured in human erythrocytes obtained from the Stanford Blood Center at 2% hematocrit in RPMI-1640 supplemented with 25 mM HEPES, 50 mg/L hypoxanthine, 2.42 mM sodium bicarbonate and 4.31 mg/ml Albumax (Invitrogen) at 37°C in 5% CO_2_ and 1% O_2_. Parasite growth assays were initially performed using fresh erythrocytes and then repeated after cryopreservation and thawing. Schizont-stage parasites were isolated using a MACS magnet (Miltenyi) and added at 0.5% initial parasitemia to previously cryopreserved wild-type or E756del heterozygous erythrocytes that had been washed and resuspended in complete RPMI with bicarbonate and Albumax as above. Assays were performed at 0.5% hematocrit in a volume of 100 μl per well in 96-well plates. For drug treatments, erythrocytes were incubated in the indicated concentrations of Yoda-1 (Sigma) for three hours at 1% hematocrit and washed in complete RPMI three times before adding to assay plates. Parasitemias were determined on day 0, day 1 (24 hours) and day 3 (72 hours) by staining with SYBR Green 1 nucleic acid stain (Invitrogen, ThermoFisher Scientific, Eugene, OR, USA) at 1:2000 dilution in PBS/0.3% BSA for 20 minutes, followed by flow cytometry analysis on a MACSQuant flow cytometer (Miltenyi).

For each genetic background (WT versus E756del heterozygote), assays were performed in technical duplicates using seven biological replicates (N=7) from unrelated donors. The percent parasitemia relative to control was determined for each biological replicate by normalizing the parasitemia on day 3 at each drug concentration relative to that in the absence of drug. Then, the mean of normalized day 3 parasitemia and S.E.M. was calculated for each genetic background at each drug concentration. Dose response inhibition curves (log inhibitor vs response) were generated in PRISM 8 Version 8.0.2 (159) and the top of the curves were constrained to equal 100 to calculate IC50 for Yoda-1. Log IC50 of the two curves were compared statistically using the extra-sum-of-squares F test.

## Results

### The PIEZO1 E756del variant is associated with protection from severe malaria

To determine if the *PIEZO1* E756del allele influences malaria susceptibility, we collected 542 DNA samples from three well-characterized malaria study cohorts from Gabon (KUN *et al.* 1998; KALMBACH *et al.* 2006; KREMSNER *et al.* 2009). Out of the 542 samples, 8 were excluded for HbSS or HbSC genotypes to minimize confounding effects of severe hematologic disease. A further 88 were excluded due to missing values in sex (n=78) and failure to amplify *PIEZO1* E756del (n=10). Therefore, our analytic cohort consisted of 446 samples. All samples were from Gabonese children between the ages of 4-140 months; 193 (43%) participants had mild malaria (controls) and 253 (57%) had severe malaria (cases). Cases of severe malaria had severe anemia, hyperparasitemia, signs of cerebral malaria, hypoglycemia and/or respiratory distress in addition to microscopically-confirmed parasitemia. Controls with mild malaria had parasitemia and fever with absence of severe signs. Their baseline demographics are summarized in Table 1. While cases and controls were similar in terms of sex (45% vs 46% male, respectively), those with severe malaria on average were younger and had higher parasite densities. Differences by study are also summarized (Table S1).

**Table 1.** Baseline demographics

| Baseline characteristic | Overall<br>(n=446) | Malaria status |  | P-value |
| --- | --- | --- | --- | --- |
|  |  | Mild<br>(n=193) | Severe<br>(n=253) |  |
| Sex, n (%) |  |  |  |  |
| Male | 201 (45%) | 88 (46%) | 113 (45%) | 0.92 |
| Female | 245 (55%) | 105 (54%) | 140 (55%) |  |
| Age in months, median (range) | 35 (4-140) | 43 (8-140) | 29 (4-133) | <0.001 |
| Parasite density (parasite/μL)<sup>1</sup>, median (range) | 35,000 (20-1,544,880) | 15,000 (588-434,074) | 108,518 (20-1,544,880) | <0.001 |
| Study, n (%) |  |  |  |  |
| *Kremsner et al.* | 195 (44%) | 49 (25%) | 146 (58%) | <0.001 |
| *Kun et al.* | 195 (44%) | 98 (51%) | 97 (38%) |  |
| *Kalmbach et al.* | 56 (12%) | 46 (24%) | 10 (4%) |  |

Continuous and categorical variables were compared across malaria status using the Wilcoxon rank sum test and Fisher's exact test, respectively. <sup>1</sup>n=44 excluded due to zero values.

The *PIEZO1* E756del variant was common among the study population, with a heterozygote prevalence of 36% in controls and 23% in cases (minor allele frequency=0.19). The variant was in Hardy-Weinberg equilibrium in the controls but not in the cases. We used a logistic regression model to predict the probability of severe malaria as a function of *PIEZO1* genotype, HbAS status, age, sex and study for each participant. The odds of severe malaria in those with the heterozygous E756del genotype (WT/DEL) were half the odds of those with the homozygous wild-type genotype (OR 0·50, 95% CI 0·31-0·81; p=0·005), suggesting that having one copy of *PIEZO1* E756del is protective against severe disease (Table 2 and Table S2). Homozygous wild-type subjects at E756 (WT/WT) had a significantly higher predicted probability of severe malaria compared to E756del heterozygous (WT/DEL) individuals (median 67% vs. 46%, p=0·01) (Figure 1A). These results suggest that the E756del variant confers protection for heterozygotes even as they age and develop increased adaptive immunity to malaria. In contrast, participants homozygous mutant for E756del (DEL/DEL) had a high probability of severe malaria compared to heterozygotes (p=0.02), suggesting that harboring two copies of this mutant allele negates the protective effect observed in carriers (Figure 1A). While participants with the homozygous mutant genotype (DEL/DEL) had over twice the odds of severe malaria compared to those who were homozygous wild-type, this association was not statistically significant, likely due to the low overall frequency of the DEL/DEL genotype (OR 2·26, 95% CI 0·82-6·94, p=0·28) (Table 2 and Figure 1A).

**Table 2.**
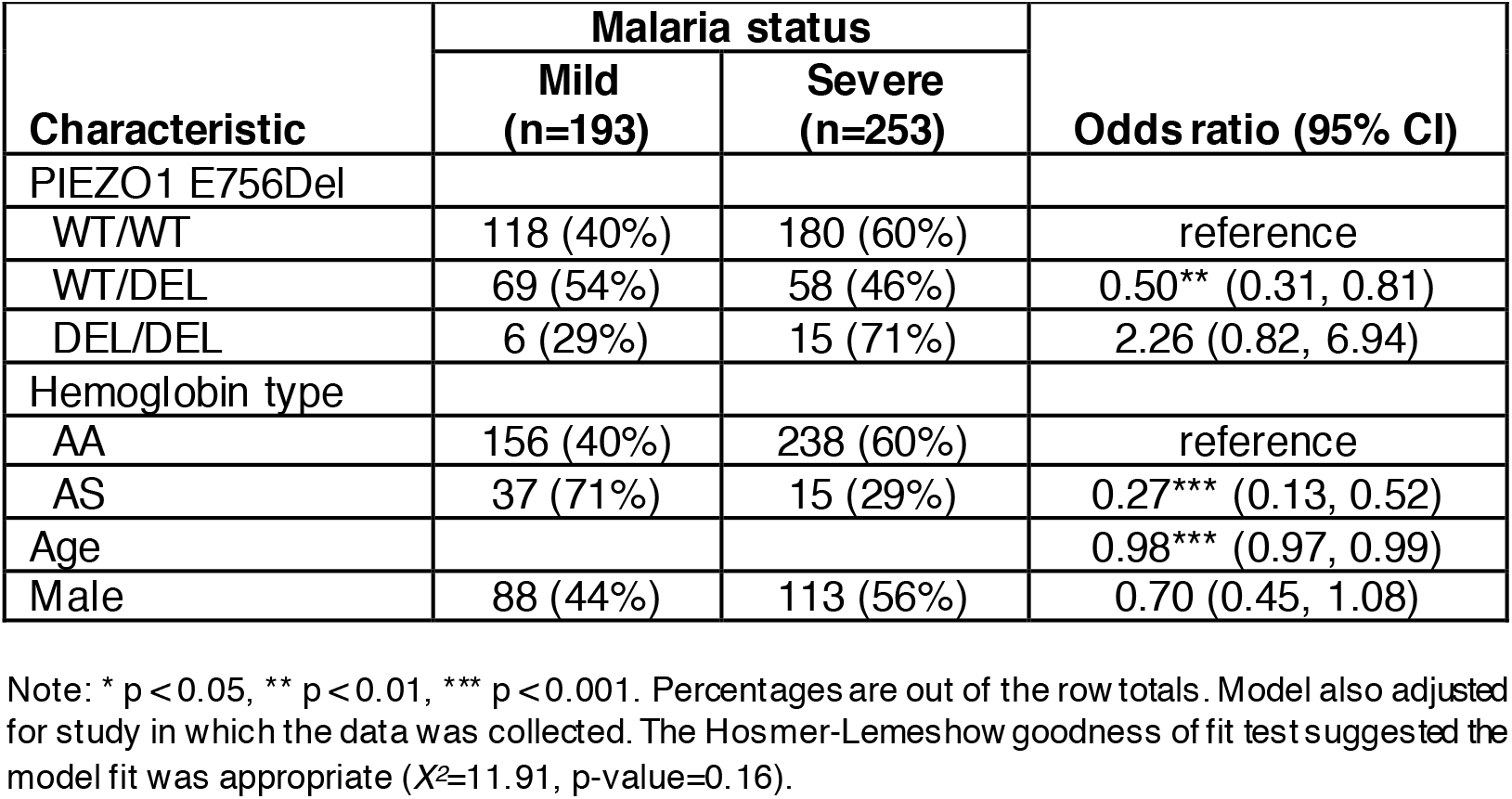
Associations with malaria severity

**Figure 1.**
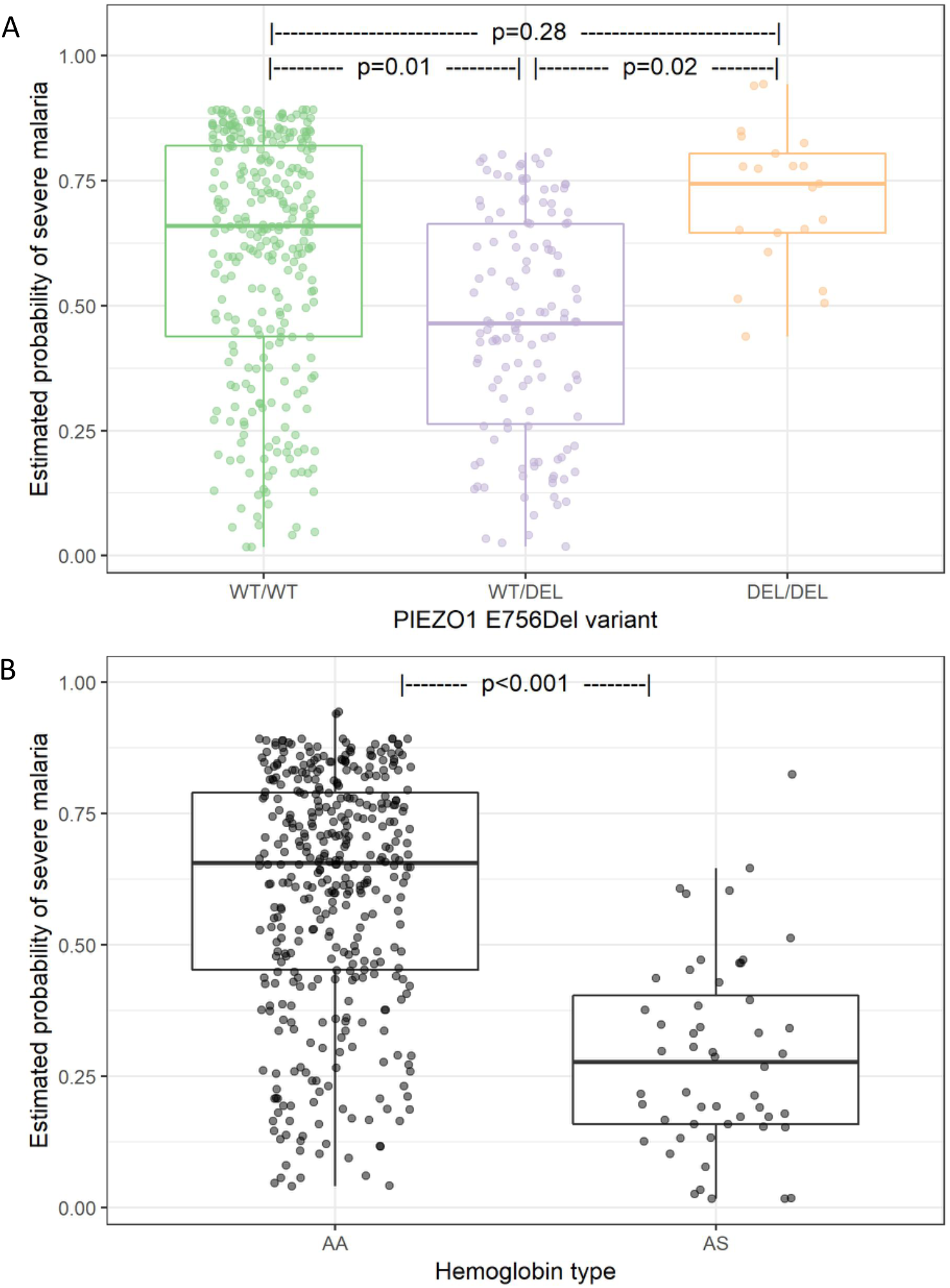
Association of malaria severity with *PIEZO1* and hemoglobin genotypes. Estimated probability of severe malaria extracted from the model presented in Table 2 by (**A**) *PIEZO1* E756 genotype and (**B**) hemoglobin beta genotype. P-values for pairwise differences in *PIEZO1* were calculated using Tukey’s HSD post hoc test. Model adjusted for age, sex, and study.

### Association between HbAS and protection from severe malaria

As expected, we also found that the sickle cell trait polymorphism (HbAS) was significantly associated with mild malaria in our study population (HbAS vs HbAA, OR 0·27, 95% CI 0·13-0·52; p<0·001) (Table 2). Children with HbAS had a significantly lower predicted probability of severe malaria compared to children with normal hemoglobin (HbAA) (28% vs 66%; p<0·001) (Figure 1B), an effect approximately twice as strong as that of *PIEZO1* E756del in our study population. This result is consistent with published literature on the protective effect of the sickle cell trait for severe malaria (MANGANO *et al.* 2015; UYOGA *et al.* 2019).

### Interplay between PIEZO1 E756del and HbS on malaria severity

Because of the high frequencies of both the *PIEZO1* E756del variant and HbAS in the study population, we sought to assess whether HbAS confounds the association found for E756del heterozygotes and malaria susceptibility by refitting the main effects model with the addition of an interaction term for HbAS and E756del. For subjects with HbAA, the E756del homozygous wild-type genotype (WT/WT) was associated with a significantly higher probability of severe malaria compared to the E756 heterozygous genotype (WT/DEL) (OR 2·09, 95% CI 1·28-3·41; p=0·003, Figure 2). In contrast, in subjects with HbAS, heterozygosity for E756del did not alter susceptibility to severe malaria compared to E756 wild-type (p=0·97, Figure 2 and Table S3). These results suggest that the heterozygous *PIEZO1* E756del genotype reduces the risk of severe malaria in individuals with normal hemoglobin, but that effect is masked by the strong protective effect of HbAS in those with sickle cell trait.

**Figure 2.**
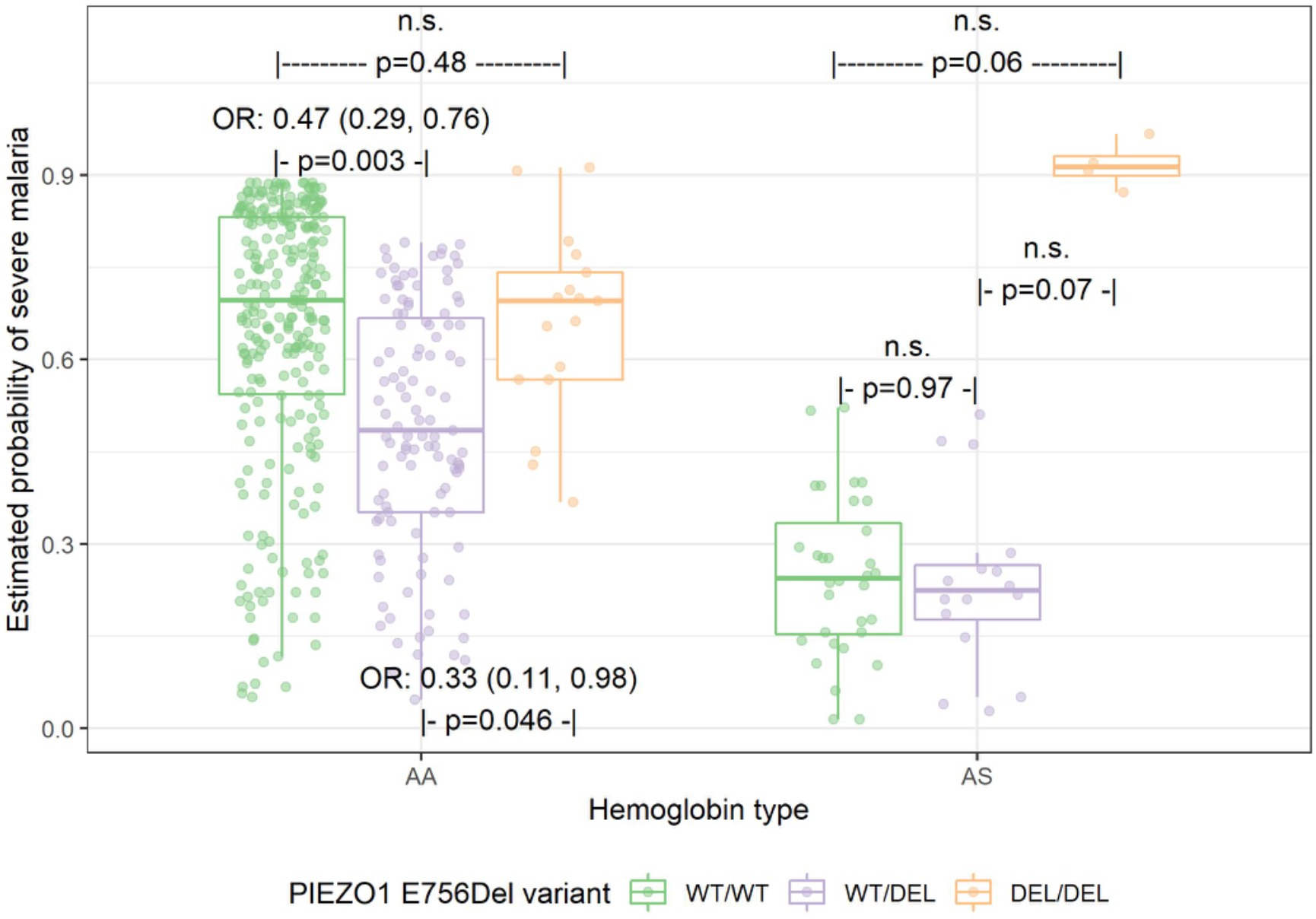
Association between *PIEZO1* and malaria severity by hemoglobin type. n.s. = not statistically significant. A Bayesian logistic regression model was fit to malaria severity status as a function of hemoglobin type, *PIEZO1*, and their interaction. Overall interaction effect of p=0.08. P-values for pairwise differences between *PIEZO1* within each hemoglobin type were calculated using Tukey’s HSD post hoc test. Model also adjusted for age, sex, and study. The Hosmer-Lemeshow goodness of fit test suggested the model fit was appropriate (*X^2^*=6.03, p-value=0.64).

For subjects with HbAA, the odds of severe malaria in those with the E756del heterozygous mutation (WT/DEL) were one third the odds of those with the homozygous mutation (OR 0·33, 95% CI 0·11-0·98; p=0·046, Figure 2). While this association was also apparent within individuals with HbAS, a difference in the risk of severe malaria failed to reach statistical significance between the two mutations (p=0·07). This result may suggest that the homozygous mutant E756del genotype alters RBC physiology in a way that abolishes the protective effect of HbAS, and may act with it synergistically to generate an unfavorable environment for controlling malaria infection. However, these results would need to be replicated in larger studies.

### Analysis of other proximal PIEZO1 polymorphisms

In addition to E756del, we identified seven other single nucleotide polymorphisms and insertion-deletions within the ~160 bp region we sequenced, which ranged in frequency from 0.4% to 7.9% (Table S4). Even though these mutations are in close proximity to E756del, none of them were significantly associated with severe malaria individually. To determine whether any of these *PIEZO1* variants act synergistically with hemoglobin type, sex or age to influence malaria susceptibility in the absence of E756del, we used classification trees (CART) (Figure 3A). For subjects with HbAA, when E756del was excluded, heterozygosity for E750Q appeared protective in younger children, whereas heterozygosity for Q749del was associated with an increased probability of severe malaria in older children. However, correlation testing showed E750Q and Q749del were significantly correlated with E756del, suggesting that these two variants may be serving as a proxy for E756del in predicting malaria severity (results not shown). This hypothesis is further strengthened when the E756del polymorphism is included in the analysis, as no other *PIEZO1* variants were predictive of disease severity in the presence of E756del (Figure 3B). Together, these results suggest that E756del is the causal variant in this region, and while two of the polymorphisms appeared to predict malaria severity, these may be serving as surrogates for E756del.

**Figure 3.**
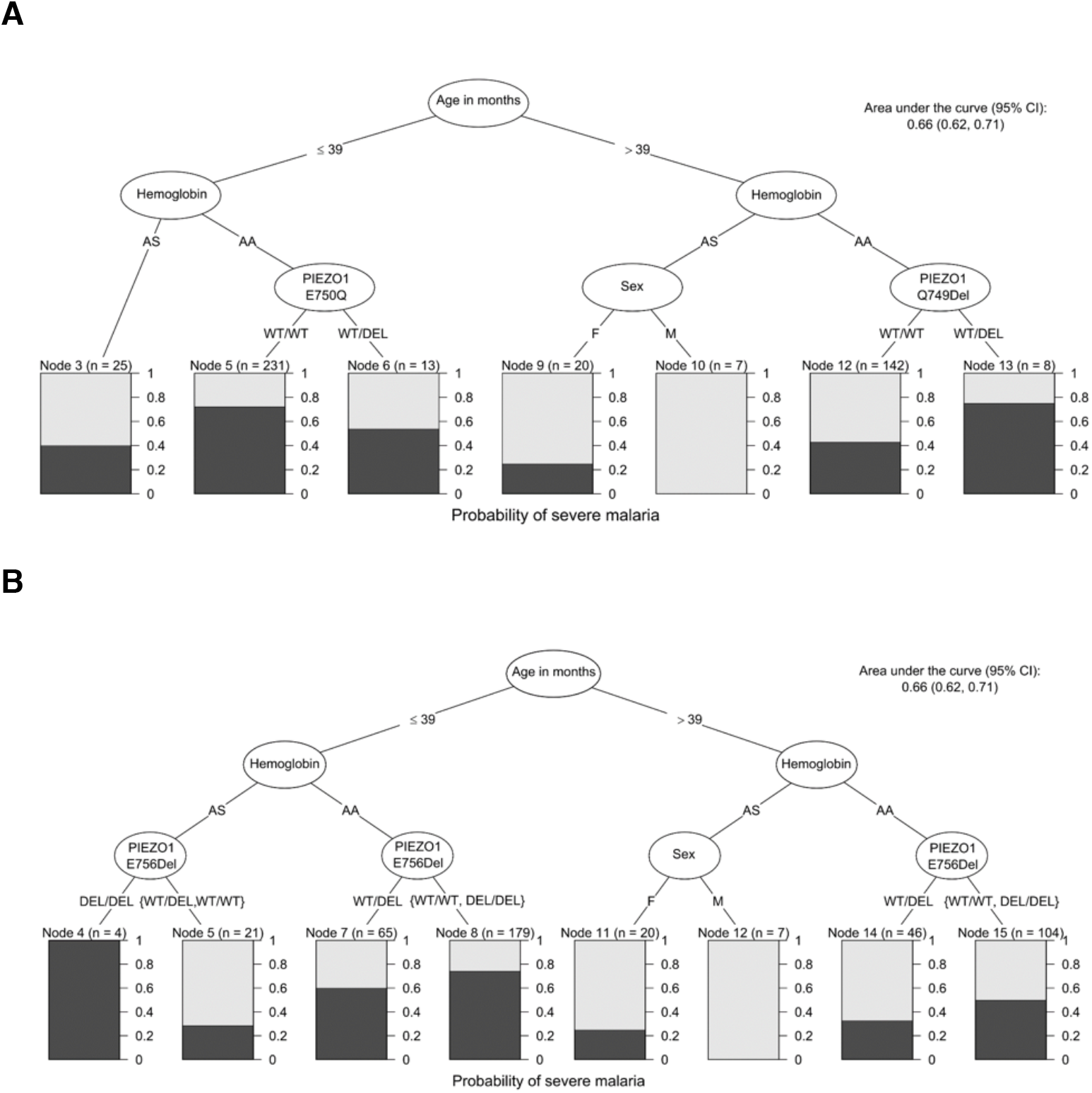
Classification trees predicting severe malaria with various possible predictors included. (A) Possible predictors: all *PIEZO1* variants except for E756Del, hemoglobin type, sex, and age. (B) Possible predictors: all *PIEZO1* variants, hemoglobin type, sex, and age. The higher the black bars, the higher the likelihood of severe malaria.

### Effect of PIEZO1 E756del on in vitro growth of P. falciparum

Previous work suggested that human red blood cells heterozygous for the *PIEZO1* E756del mutation are less supportive of *P. falciparum in vitro* growth than wild-type red blood cells (MA *et al.* 2018). To further investigate whether impaired intracellular growth could be a mechanism by which E756del protects against severe malaria, we identified local donors of African descent who are wild-type or heterozygous at *PIEZO1* E756del and used their red blood cells in *P. falciparum* growth assays. To our surprise, we found that *P. falciparum* replicated equally well in the wild-type versus E756del heterozygous erythrocyte samples (p=0.37, Figure 4A). Similar trends were observed when the cells were used freshly or after cryopreservation. These results suggest that while the E756del mutation may confer gain-of-function kinetics (MA *et al.* 2018), at least in heterozygous RBCs, the phenotype is not strong enough to impair intracellular growth of *P. falciparum in vitro*.

**Figure 4.**
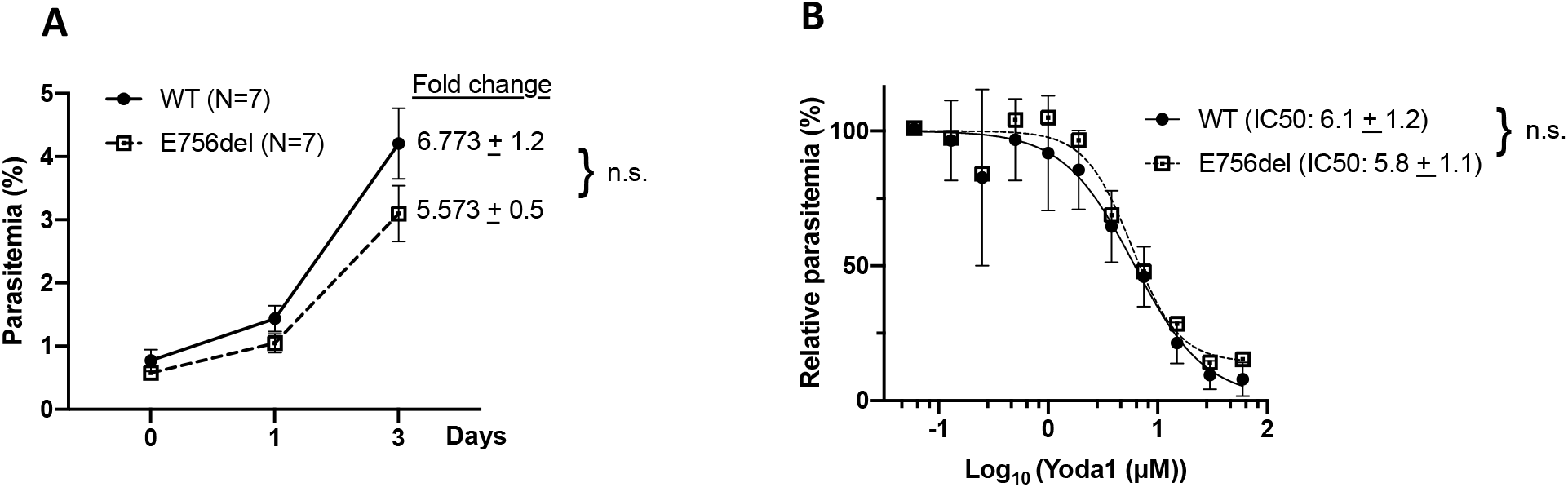
*P. falciparum* growth is preserved in E756del erythrocytes but a small molecule agonist of PIEZO1 has antimalarial activity. (**A**) Growth curves for *P. falciparum* strain 3D7 in WT or *PIEZO1* E765del heterozygous erythrocytes. Each point represents mean raw parasitemia of N=7 donors for each genetic background, and error bars represent SEM. Each sample was run with two technical replicates. “Fold change” indicates the average fold difference (+SEM) in parasitemia on day 3 relative to day 0. n.s., not significant, p-value=0.3725. (**B**) *P. falciparum* strain 3D7 parasitemia as a function of Yoda1 concentration, as determined by flow cytometry after 3 days of drug treatment. The relative parasitemia was calculated for each biological replicate (N=7 for each genetic background) by normalizing the day 3 parasitemia at each drug concentration (two technical replicates) relative to the parasitemia in the absence of drug. Yoda1 IC50 ± SEM (*μ*M) for WT or E765del erythrocytes are shown in parenthesis. n.s., not significant, p-value=0.4639. SEM=Standard error of mean.

To determine if chemical activation of the PIEZO1 ion channel could inhibit intracellular growth of *P. falciparum* in RBCs, we used the small synthetic molecule Yoda1, which was recently identified as a specific agonist of both human and mouse PIEZO1 (SYEDA *et al.* 2015). We observed a dose-dependent effect on *P. falciparum* growth in both wild-type and E756del RBCs, with an IC50 of ~6 μM (Figure 4B). This IC50 concentration is similar to that required for Yoda1 to induce PIEZO1-specific Ca^2+^ responses in HEK cells or mouse RBCs (CAHALAN *et al.* 2015; SYEDA *et al.* 2015). These results demonstrate that Yoda1 has antimalarial activity and point to a critical role for erythrocyte PIEZO1 in promoting the development of *P. falciparum* in red blood cells. The observation that the antimalarial potency of Yoda1 is similar in the wild-type and E756del genetic backgrounds provides additional evidence that the E756del mutation likely confers only a mild gain-of-function phenotype in heterozygous RBCs.

## Discussion

In this work, we sought to determine the clinical relevance of a recent study demonstrating a survival benefit for *Plasmodium*-infected mice carrying a *PIEZO1* gain-of-function allele (MA *et al.* 2018). In the *PIEZO1*GOF mouse model, the RBCs were dehydrated, intracellular parasite growth was impaired, and the animals were protected from developing cerebral malaria. We assessed the human *PIEZO1* mutation E756del, which was identified as a gain-of-function allele based on its prolonged inactivation time constant after mechanical stimulation in HEK cells (MA *et al.* 2018). Using samples from Gabonese patients with severe or uncomplicated malaria, we found that *PIEZO1* E756del is significantly associated with protection against severe malaria. Children heterozygous for E756del had a significantly lower predicted probability of severe malaria compared to those with wild-type *PIEZO1*, even when controlling for age, sex, and HbAS status. As E756del is present at high frequencies in African populations (MA *et al.* 2018), these results suggest that it may be a major determinant of innate resistance to severe malaria. Whether this advantage is extended to all severe malaria sub-phenotypes remains to be investigated in the context of a larger study.

Given the strong protective effect of *PIEZO1* E756del on severe malaria in our study, its absence from GWAS candidate lists is particularly notable. The sensitivity of GWAS studies for malaria in Africa are limited by several factors, including weak linkage disequilibrium and population stratification (JALLOW *et al.* 2009), as well as technical limitations in assessing complex or repetitive regions via microarrays or deep sequencing. The *PIEZO1* E756del mutation is a three nucleotide deletion within a region of short tandem repeats, making it difficult to detect by high-throughput methods. In our study, accurate genotyping of this locus required Sanger sequencing and manual alignments. These studies on *PIEZO1* highlight the benefit of using a combination of sequencing methods for the identification of host factors that may influence susceptibility to malaria.

The finding that E756del is protective against severe malaria only in heterozygotes is reminiscent of the protective effect of HbAS, where heterozygotes with sickle cell trait are protected from malaria-related mortality but homozygotes with sickle cell disease are not (AMBE *et al.* 2001; AIDOO *et al.* 2002; KOMBA *et al.* 2009). Although GOF mutations in *PIEZO1* are a recognized cause of hereditary xerocytosis (GLOGOWSKA *et al.* 2017), to our knowledge the E756del allele has not been implicated in any hematologic disease (MA *et al.* 2018). If E756del is truly a GOF allele it seems likely that homozygous mutant cells would be dehydrated and display altered permeability, predisposing to hemolytic anemia and severe symptoms upon infection with *P. falciparum*. Given the high allele frequency of E756del in our Gabonese study population (19%), understanding how homozygosity for this mutation affects PIEZO1 channel function, RBC hydration status and *in vitro* susceptibility to *P. falciparum* are important open questions.

We did not observe any influence of co-inheritance of HbAS and E756del heterozygosity on malaria susceptibility, suggesting that having one E756del allele does not contribute any additional effect to the already powerful protection conferred by HbAS. In contrast, subjects with HbAS who are homozygous mutant for E756del had an extremely high predicted probability of severe malaria. While this association was not statistically significant, it highlights the need for further research on *PIEZO1* E756del, as it suggests that the protective effect of HbAS on malaria can be subverted by structural or permeability abnormalities found in homozygous E756del RBCs. Therefore, future studies would need to further validate these findings. A recent analysis of associations between E756del and clinical parameters in sickle cell disease (SCD) patients showed that the *PIEZO1* E756del variant was associated with increased RBC density and dehydration in HbSS cells, providing an additional example of a potential phenotypic interplay between these genes (ILBOUDO *et al.* 2018).

The hypothesis that *PIEZO1* may influence malaria susceptibility arose from the observation that *P. falciparum* parasites grow poorly in dehydrated red blood cells and the knowledge that hereditary xerocytosis, a disorder characterized by red blood cell dehydration, is often caused by GOF mutations in *PIEZO1* (TIFFERT *et al.* 2005; ANDOLFO *et al.* 2013; ARCHER *et al.* 2014). Although our results show that *PIEZO1* E756del is protective against severe malaria in heterozygotes, the mechanism of protection is not yet known. In the mouse model, expression of a *PIEZO1* GOF allele in the hematopoietic lineage was protective against cerebral malaria and appeared to impair growth of *P. berghei* ANKA parasites, at least during the initial phase of infection (MA *et al.* 2018). The same study suggested that *P. falciparum* growth was impaired in human RBCs carrying an E756del allele (MA *et al.* 2018). However, in our experiments we did not observe a significant *P. falciparum* growth defect in *PIEZO1* E756del heterozygous RBCs. *P. falciparum* grew equally well in wild-type versus E756del-carrying RBCs, regardless of whether the cells were fresh or had been previously cryopreserved. These results demonstrate that heterozygosity for *PIEZO1* E756del does not in itself hinder *P. falciparum* invasion or growth in red blood cells. While the reason for the discrepant findings is not clear, it is possible that time from donation, processing variability and/or other technical issues confounded the conclusions of the previous study.

Our studies indicate that while *P. falciparum* can grow normally in human RBCs carrying a *PIEZO1* E756del allele, the PIEZO1 agonist Yoda1 had potent antimalarial activity. Presumably the changes in ionic permeability induced by Yoda1 create an inhospitable environment for normal parasite development. Since the drug assay results demonstrate that strong activation of PIEZO1 in RBCs can inhibit *P. falciparum* growth, one explanation for the lack of a phenotype in untreated E756del heterozygous RBCs could be that E756del confers only a mild channel defect. Indeed, out of two candidates and one established *PIEZO1* GOF mutation tested, E756del displayed the shortest channel inactivation time (MA *et al.* 2018). As the experiments measuring channel kinetics were performed using ectopic expression in HEK cells, it is possible that the channel phenotype of E756del in heterozygous RBCs is even more mild. This conclusion is further supported by our findings showing that wild-type and E756del RBCs were equally sensitive to Yoda1’s antimalarial activity. Previous work in mice showed that Yoda1 is specific for PIEZO1, as it induced cation permeability and cellular dehydration in wild-type but not PIEZO1-null mouse RBCs (CAHALAN *et al.* 2015). Bioinformatic analyses aimed at identifying mechanosensitive channels in pathogenic protozoa did not identify any homologues of *PIEZO1* in Apicomplexan parasites, making it unlikely that Yoda1 could be acting on a parasite-encoded target (PROLE AND TAYLOR 2013). In our experiments, we further minimized this possibility by pre-incubating the RBCs in Yoda1 for several hours and then washing the cells extensively before infecting with parasites.

Given that *in vitro* growth of *P. falciparum* in RBCs is unaffected by PIEZO1 E756del, how might this common polymorphism protect from severe malaria? As with many life-threatening systemic infections, the symptoms of severe malaria are in large part caused by the body’s extreme response to overwhelming infection by a microorganism. Additionally, the adhesive properties of *P. falciparum*-infected RBCs enable them to sequester in the deep microvasculature and adhere to uninfected RBCs, exacerbating organ dysfunction. As *PIEZO1* is expressed in many tissues including the vascular endothelium and immune cells, it is possible that dysregulation of its channel activity has effects on vascular tone, cell permeability and/or signaling during malaria infection that are independent of any effects on RBCs. Alternatively, E756del may alter *P. falciparum* virulence properties rather than intracellular growth, such as the ability of the parasite to export cytoadherence proteins to the RBC plasma membrane. This type of mechanism has previously been proposed to explain the malaria-protective nature of the hemoglobin C mutation (FAIRHURST *et al.* 2005).

Our findings show a significant association between *PIEZO1* E756del and protection from severe *P. falciparum* malaria. Together with the previous elegant mechanistic work on mouse *PIEZO1* GOF mutations and *P. berghei* ANKA, these data firmly establish PIEZO1 as an important host susceptibility factor for malaria. Though the list of innate susceptibility determinants for severe malaria includes a range of factors such as hemoglobin variants, enzymes, membrane proteins and immune-related molecules (APINJOH *et al.* 2013; NGUETSE AND EGAN 2019), PIEZO1 is unique because of its potential to influence *P. falciparum* pathogenesis on multiple levels (including its asexual replication in RBCs and ability to mediate cerebral malaria), and because it is considered a druggable molecule (SYEDA *et al.* 2015). The demonstration that chemical activation of RBC PIEZO1 impairs *P. falciparum* growth *in vitro,* combined with the discovery that the E756del allele is associated with protection from severe malaria in patients, suggests that PIEZO1 has potential as a compelling target for a host-directed therapy for malaria.

## Author contributions

CNN, experimental design, acquisition, analysis and interpretation of data, and writing the manuscript; NP, analysis and interpretation of data and editing the manuscript; BS, acquisition and analysis of data and editing the manuscript; ERE, conception and experimental design, sample acquisition, data analysis and interpretation and editing the manuscript; PGK, TPV, sample acquisition, experimental design, and editing the manuscript; ESE, conception and experimental design, analysis and interpretation of data, provision of resources and supervision, and writing the manuscript. All authors read and approved the final manuscript.

## Acknowledgments

We thank the staff of Albert Schweitzer Hospital in Lambaréné, and the Centre Hospitalier de Libreville in Gabon for their assistance with samples and data collection; and the study patients and their parents for consenting to participate in those studies. We thank Frans Kuypers, Ardem Patapoutian, Elizabeth Winzeler, and members of the Boothroyd, Petrov, Yeh and Egan labs for helpful discussions. We thank Carrie Lin, Nick Bondy and Tuya Yokoyama for technical assistance. Local blood samples were drawn at the Stanford Clinical and Translational Research Unit, which is supported by CTSA Grant # UL1 TR001085 CNN received support from the Stanford Maternal and Child Health Research Institute, and ESE was funded through awards from the Doris Duke Charitable Foundation (#2016098), NIH (DP2HL13718601), the Stanford Maternal Child Health Research Institute, and is a Tashia and John Morgridge Endowed Faculty Scholar in Pediatric Translational Medicine and a Baxter Faculty Scholar.

