## Supplemental files for "A common polymorphism in the druggable ion channel *PIEZO1* is associated with protection from severe malaria"

### Supplemental Tables and Figures

**Table S1.** Baseline demographics by study

| Baseline characteristic | Overall (n=446) | Study *Kremsner* (n=195) | *Kun* (n=195) | *Kalmbach* (n=56) |
|---|---|---|---|---|
| Sex, n (%) | | | | |
| Male | 201 (45%) | 104 (53%) | 74 (38%) | 23 (41%) |
| Female | 245 (55%) | 91 (47%) | 121 (62%) | 33 (59%) |
| Age in months, median (range) | 35 (4, 140) | 25 (12, 103) | 41 (8, 140) | 36 (4, 120) |
| Parasite density (parasite/µL)[1], geometric mean (95% confidence interval) | 34,664.9 (34,649.5, 34,680.3) | 26,552.0 (26,536, 26,568.1) | 42,643.6 (42,626.9, 42,660.3) | 34,581.9 (34,572.7, 34,591.0) |
| Severe malaria, n (%) | 253 (57%) | 146 (75%) | 97 (50%) | 10 (18%) |

Continuous and categorical variables were compared across malaria status using the Wilcoxon rank sum test and Fisher's exact test, respectively. [1]n=44 excluded due to zero values.

**Table S2.** Associations with malaria severity – multiple imputation (n=534)†

| Characteristic | Mild malaria | Severe malaria | Odds ratio (95% CI) |
|---|---|---|---|
| PIEZO1 E756Del | | | |
| WT/WT | 63% | 71% | reference |
| WT/DEL | 34% | 23% | 0.54** (0.35, 0.84) |
| DEL/DEL | 3% | 6% | 2.11 (0.82, 5.42) |
| Hemoglobin | | | |
| AA | 80% | 95% | reference |
| AS | 20% | 5% | 0.22*** (0.11, 0.42) |
| Age | 40 (4, 140) | 28 (4, 133) | 0.98*** (0.97, 0.99) |
| Male | 48% | 44% | 0.69 (0.44, 1.09) |

Note: * $p < 0.05$, ** $p < 0.01$, *** $p < 0.001$. Model also adjusted for study in which the data was collected. †Multiple imputation resulted in 5 imputed data sets, thus only percentages by malaria status are shown for the pooled data.

**Table S3.** Associations with malaria severity including hemoglobin type: multiple imputation (n=534) †

| Characteristic | Odds ratio (95% CI) |
|---|---|
| PIEZO1 E756Del (ref=WT/WT) | |
| WT/DEL | 0.51** (0.33, 0.80) |
| DEL/DEL | 1.38 (0.53, 3.59) |
| Hemoglobin AS (ref=AA) | 0.15*** (0.07, 0.34) |
| PIEZO1 and Hemoglobin interaction | |
| WT/DEL * AS | 2.26 (0.57, 8.92) |
| DEL/DEL * AS | 19.73 (0.82, 476.59) |
| Age | 0.98*** (0.97, 0.99) |
| Male | 0.70 (0.45, 1.10) |

Note: * $p < 0.05$, ** $p < 0.01$, *** $p < 0.001$. Bayesian logistic model also adjusted for study in which the data was collected. Interaction $p = 0.11$. †Multiple imputation resulted in 5 imputed data sets, thus only percentages by malaria status are shown for the pooled data.

**Table S4.** Associations with malaria severity among other PIEZO1 variants (n=532[1])

| PIEZO1 variant | Malaria status: Mild (n=252) | Malaria status: Severe (n=280) | Odds ratio (95% CI) |
|---|---|---|---|
| **E756EE** | | | |
| WT/WT | 248 (98.4%) | 277 (98.9%) | 1.49 (0.33, 7.62) |
| WT/DEL | 4 (1.6%) | 3 (1.1%) | reference |
| **E755-E756Del[2]** | | | |
| WT/WT | 252 (100%) | 274 (97.9%) | 0.07 (0.00, 1.23) |
| WT/DEL | 0 | 6 (2.1%) | reference |
| **E750Q** | | | |
| WT/WT | 236 (93.7%) | 264 (94.3%) | 1.12 (0.54, 2.30) |
| WT/DEL | 16 (6.3%) | 16 (5.7%) | reference |
| **Q749-E750Del[2]** | | | |
| WT/WT | 252 (100%) | 279 (99.6%) | 0.32 (0.01, 8.07) |
| WT/DEL | 0 | 1 (0.4%) | reference |
| **Q749Del** | | | |
| WT/WT | 240 (95.2%) | 255 (91.1%) | 0.51 (0.24, 1.02) |
| WT/DEL | 12 (4.8%) | 22 (7.9%) | reference |
| DEL/DEL | 0 | 3 (1.1%) | |
| **Q749E** | | | |
| WT/WT | 251 (99.6%) | 275 (98.2%) | 0.22 (0.01, 1.37) |
| WT/DEL | 1 (0.4%) | 5 (1.8%) | reference |
| **E739Q[2]** | | | |
| WT/WT | 250 (99.2%) | 280 (100%) | 6.19 (0.31, 124.48) |
| WT/DEL | 2 (0.8%) | 0 | reference |
| **At least 1 non-WT** | | | |
| WT/WT | 217 (86.1%) | 227 (81.1%) | 1.45 (0.91, 2.32) |
| WT/DEL | 35 (13.9%) | 53 (18.9%) | reference |

[1] n=10 excluded due to missing values.

[2] Due to estimation issues, fit using Bayesian logistic regression.
